# Predicting Binding Affinities for the Binding Domain of Hyperpolarization-Activated Cyclic Nucleotide-Gated Channel Isoforms Using Free-Energy Perturbation

**DOI:** 10.64898/2026.03.04.709733

**Authors:** Matthew Brownd, Stephanie Sauve, Hope Woods, Mahmoud Moradi

## Abstract

Hyperpolarization-activated cyclic nucleotide-gated (HCN) channels are are a family of voltage-gated, cyclic-nucleotide modulated *Na*^+^/*K*^+^ channels that regulate spontaneous rhythmic electrical activity in both the heart and the brain. Understanding differences in the responsiveness to cyclic adenosine monophosphate (cAMP) modulation between HCN isoforms would offer insight into the specific binding interactions that drive channel activation. Using all-atom molecular dynamics (MD) simulations and the free-energy perturbation (FEP) approach, we determined the absolute binding free energy of cAMP to the the cyclicnucleotide-binding domain (CNBD) of HCN isoforms 1-4. By studying the free-energy of ligand binding to the various isoforms of HCN, our study advances the understanding of HCN channel activation and modulation mechanisms. Overall, our work offers insight into explaining differences in channel sensitivity across the isoforms of HCN.

## Introduction

Hyperpolarization-activated cyclic nucleotide-gated (HCN) channels are a family of voltagegated, cyclic-nucleotide modulated *Na*^+^/*K*^+^ channels that are expressed in the heart and central nervous system. ^1–3^ Referred to as “pacemaker” channels, they function as regulators of spontaneous rhythmic electrical activity in both the heart and the brain. ^4^ The dysfunction of HCN channels has been linked to arrhythmia and a wide variety of neurological disorders, including epilepsy, Parkinson’s disease, and generalized seizures. ^5,6^ HCN channels have also been shown to play a role in the perception of pain, making them a potential target for analgesics.^7^ All of these factors would seem to indicate that HCN channels are an attractive target for therapeutic drug design, however, as of 2020 only a single drug targeting HCN channels has been approved by the FDA - Ivabradine, which is used to treat angina pectoris and chronic heart failure.^8^ One of the greatest barriers hindering progress towards targeting the HCN channel family is the fact that their conformational dynamics and the precise mechanistic details of their dual activation by both voltage and the modulatory ligand cyclic adenosine monophosphate (cAMP) are still poorly understood.^9^

The binding of cAMP to the cyclic nucleotide-binding domain (CNBD) of HCN channels inside the cell has shown to have several effects on the channels, including a slight depolarizing shift in the voltage threshold necessary for channel activation,^10^ kinetically favoring the open channel conformation,^11^ and promoting channel tetramerization,^12^ all of which contribute to an overall improved channel conductance. However, the binding of cAMP in the absence of a hyperpolarizing membrane potential is still not sufficient to induce opening of the channel. ^1,3^ According to truncation studies which were performed previously, ^4^ the purpose of the CNBD seems to be to inhibit activation of the core transmembrane helix bundle which serves as the channel pore, and the binding of cAMP acts to relieve this inhibition.^9^ Differences in the activation gating and responsiveness to cAMP modulation between HCN isoforms would therefore be the result of variation in the amino acid residue sequence/conformation of the CNBD, leading to differences in the effective steric inhibition between the channels.^4^

Much of the current understanding surrounding the behavior and function of HCN channels has come from comparing solved structural “snapshots” of different channel states (ie. apo vs. holo, or active vs. inactive), identifying the differences, and then trying to propose a mechanism which fills in the blanks (see Lee & MacKinnon, 2017).^13^ An all-atom molecular dynamics (MD) approach, on the other hand, provides an unparalleled degree of structural detail and dynamic conformational insight by allowing regions of the conformational landscape which were previously inaccessible to be sampled and studied. Here, we conducted a series of all-atom MD simulations on various isoforms of the HCN channel family to gain deeper insights into the specific binding interactions that drive channel activation and modulation, as well as the factors contributing to differences in channel sensitivity. Using both equilibrium and non-equilibrium all-atom MD simulations, we have studied the absolute binding free energy of cAMP to the CNBDs of monomeric HCN1-4 isoforms.

In summary, our comparative MD simulations enhance the understanding of cAMP modulation across isoforms of HCN channels. In our studies, we observe similar binding behavior across the isoforms to cAMP, suggesting that the differences in channel activation across the isoforms is not merely a result of the amino acid residue sequences of the CNBD. Our study progresses the understanding of HCN channel modulation by cAMP and atomistic insights from our simulations could guide the design of functionally selective drugs targeting isoforms of this important channel family.

## Methods

### Molecular Modeling and Simulation Systems

We employed all-atom MD simulations to elucidate the binding action of cyclic nucleotides to the CNBD of monomeric HCN isoforms. To do this, we designed four simulation systems based on the CNBD of monomeric HCN1-4 proteins bound by a cAMP ligand. For HCN1, a cryogenic-electron (cryo-EM) microscopy structure of cAMP bound to HCN1 was used as an initial structure (PDB: 5U6P, 3.51 Å resolution).^13^ To isolate the CNBD region, the structure was manually truncated to residues D445-D608. For HCN2, an X-ray crystallography structure of cAMP bound to HCN2 was used as an initial structure (PDB: 3U10, 2.30 Å resolution).^12^ The structure was truncated to residues D514-D677 to isolate the CNBD region. For HCN3, an AlphaFold ^14–16^ structure (AF-Q9P1Z3-F1-v4) was obtained from Uniprot ^17^ because the structure has not yet been determined experimentally. The model was truncated to residues D398-R558 to isolate the CNBD region. The cAMP was inserted into the HCN3 structure by aligning the HCN3 model with the HCN1 model which was used as a template. For HCN4, an X-ray crystallography structure where cAMP is bound to HCN4 was used as an initial structure (PDB: 3U11, 2.50 Å resolution).^12^ The structure was truncated to residues D565-D728 to isolate the CNBD region. To ensure that all systems were of similar length and included the full D-helix, any missing terminal residues were modeled using the HCN1 structure as a template and the necessary mutations for the appropriate isoform sequence were made.

The prepared CNBD of HCN isoform systems were made using the Solution Builder module in CHARMM-GUI^18^ in which 0.15 M NaCl was included (in addition to the counterions used to neutralize the protein) to mimic physiological conditions. The simulation systems were solvated in a cubic box of TIP3P water (HCN1 systems had dimensions of 83 Å × 83 Å × 83 Å, HCN2 systems had dimensions of 84 Å × 84 Å × 84 Å, HCN3 systems had dimensions of 85 Å × 85 Å × 85 Å, and HCN4 systems had dimensions of 83 Å × 83 Å × 83 Å). The total number of atoms in the ligand-bound systems were 53,664 atoms for HCN1, 55,640 for HCN2, 57,363 for HCN3 and 53,866 for HCN4. All of the ligand-bound HCN isoform systems were simulated with ten replicates. Additionally, a system containing just the cAMP ligand solvated in TIP3P waters and 0.15 M NaCl to mimic physiological conditions was also simulated for the free energy perturbation. For this system that contained 7,410 atoms, the cubic box had dimensions of 43 Å × 43 Å × 43 Å and only one replicate was simulated. All MD simulations were performed in NAMD 2.14^19^ using the CHARMM36m force field^20^ for proteins and ions. For the cAMP ligand, the CGenFF force field was used. Energy minimization was initially applied to the HCN containing systems using the conjugate gradient algorithm for 10,000 steps. Production runs were then carried out for 100 ns for each replicate, for an aggregate simulation time of 1 *µ*s for each isoform. For the ligand only system, the system was equilibrated for 2.5 ns and the production run was carried out for 50 ns.

The equilibration phase was performed in an NVT ensemble, while the production runs were conducted in an NPT ensemble. Each simulation consisted of 2.5 ns of equilibration (25 ns for HCN3, due to the manual insertion of the ligand), followed by 10 ns of production with restraints. For the restraints, a distance restraint between “resname CMP” and “protein and within 5 of resname CMP” was used and a RMSD harmonic center restraint between “resname CMP” when aligned with “protein and alpha and sheet”, using a force constant of 5 kcal/mol/*^°^A*^2^, was used. These restraints were designed to maintain the ligand’s position relative to the *β*-jelly roll. The production time with restraints was followed by 2 ns of linearly lifting that restraint. A time step of 2 fs was utilized, and the temperature was maintained at 310 K using a Langevin thermostat. The Nosé-Hoover Langevin piston method^21^ was used to control pressure at 1 bar.

### Trajectory Analysis

Center of mass displacements were conducted on the ligand based on the center of mass relative to the ligand’s starting position. The root-mean-square deviation (RMSD) trajectory tool in VMD^22^ was employed to calculate the RMSD, with all atoms of the ligand considered for these calculations. The “lid distance” is defined as the distance between the *C*_*α*_ atoms of a particular residue in a given loop of the *β*-jelly roll and a particular residue in the turn between helices C and D. The residue pairs used were S527/I594, N596/I663, A480/I547, and N647/I714 for HCN1, HCN2, HCN3, and HCN4 respectively. For the angle analysis, the *C*_*α*_ atoms of the residues in the B and C helices were used to calculate the moment of inertia of the individual helices, and the angle between these vectors was determined. Hydrogen bond analysis was conducted using the VMD HBond plugin, ^22^ with a default cutoff distance of 3.0 Å and an angle cutoff of 20°.

### Free-Energy Perturbation

Trajectories for each isoform were chosen for alchemical free energy perturbation only if the docking pose was determined to be suitable. Here, four trajectories (six in the case of HCN3, due to the nature of the ligand insertion as well as the lack of pre-existing data) of the ligand-bound systems were chosen as having the most suitable docking poses based on low ligand center of mass displacement, low ligand RMSD, low lid distance values, and stable hydrogen bonds with key residues of interest. From each of these trajectories, conformations were saved from the production runs at 20 ns intervals (5 per system) to be used as seeds for FEP^23^ binding free energy calculations. Additionally, three replicates at the end of the simulation (50ns of production) of the solvated ligand system were used as seeds for FEP.

The FEP approach^24^ was used to calculate the relative binding free energies of cAMP to the CNBD of HCN1-4. The FEP process consisted of gradually annihilating (forward process) or returning (backward process) the cAMP ligands for the systems in question to +simulate the binding of cAMP to the HCN isoforms and obtain binding free energies. Each simulation comprised 20 windows (Δ*λ*=0.05 per window), with each window consisting of 10 ps of alchemical equilibration followed by 115 ps of data collection (2.5 ns total). Every window for a given simulation was initialized with the same initial seed conformation coming from the previously described equilibrated trajectories, but with the nucleotide already partially annihilated, allowing all windows to be run simultaneously in parallel. All FEP simulations were carried out with a timestep of 1 fs. To circumvent the endpoint problem for van der Waals interactions,^25,26^ a soft-core potential was used such that the electrostatic interactions were fully decoupled from the system by *λ*=0.5. Free energy differences were calculated using the Bennett Acceptance Ratio (BAR)^27^ method. The resulting data from FEP calculations were statistically analyzed using the VMD ^22^ ParseFEP plugin^28^ to obtain the relative binding free energies of cAMP to the CNBD of HCN1-4.

### Per-Residue Interaction Free Energies

The FEP trajectories were used to determine the relative per-residue interaction free energies for cAMP using the pairinteraction function of NAMD to calculate the potential energies between residues of interest and the transitioning nucleotide. Residues were selected for this analysis based on the existence of any hydrogen bonds which were identified in the starting structure, as well as any residues which were not fully conserved between the isoforms while rejecting any which were too far away or could not directly access the ligand.

The electric and van der Waals potential energy terms between these residues and the transitioning nucleotide were calculated every 5 ps 25 times per window, or 500 times per doublewide FEP process. We approached the alchemical transformation of the ligand as an equilibrium problem between two states, one where the ligand is fully present and one where it is fully annihilated, allowing us to calculate an equilibrium constant, K, for each window as an average of exponentials of the difference in sampled potential energies between states as shown below, where RT=0.616 kcal/mol. Since the electric and van der Waals terms scale differently in the FEP simulations, Equation 1 is used for the first ten windows, and Equation 2 is used for the last ten windows:

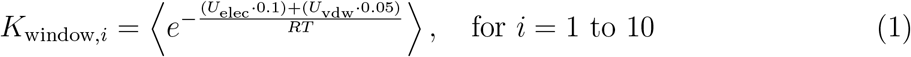

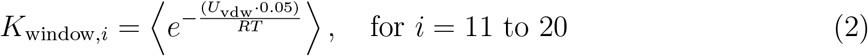

Ultimately, the relative binding interaction free energy for a given residue with respect to cAMP can be approximated as shown in Equation 3, where the output is in kcal/mol:

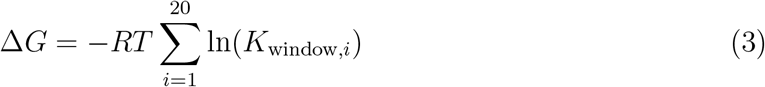

The values obtained through this analysis are used to discuss the relative importance of specific residue interactions in the binding of HCN1-4 to cAMP.

## Results and Discussion

Before conducting FEP to obtain absolute binding free-energies for cAMP binding to the CNBD of isoforms of HCN, we first identified suitable replicates to conduct FEP on for each isoform. Specific replicates were chosen to ensure that stable ligand binding was present after the restraints on the ligand were lifted for the FEP step. In the following section we first show results explaining our criteria for choosing the replicates chosen to conduct FEP on for each isoform (see Supplemental Information for the data for the replicates not chosen for FEP). In general, replicates were chosen if they had low ligand center of mass displacement, low ligand RMSD, low lid distance values, and stable hydrogen bonds with key residues of interest. In the later section, we discuss the results of our FEP approach and rank the relative binding of cAMP to the HCN isoforms. Finally, we discuss the importance of conserved arginine and glutamate residues in the CNBDs of the isoforms based on per-residue interaction free-energies.

### Choosing Suitable Replicates for FEP

For our first selection criteria, replicates were advanced to the FEP analysis stage if the ligand center of mass displacement was considered to be small, below 2 Å, as shown in Figure 1 for the replicates that were selected for FEP. A small ligand center of mass displacement indicates that the ligand was not changing its overall position in the binding pocket, further suggesting that the ligand was not moving out of the pocket and was stably bound. Additionally, this was confirmed by visually viewing the trajectories of the simulations in VMD^22^ to confirm that the ligand did not exit the binding site during the simulations qualitatively. Although the replicates chosen for HCN2 and HCN4 had lower ligand center of mass displacements compared to HCN1 and HCN3, for the majority of the simulations, all chosen replicates for all isoforms were below the 2 Å threshold.

**Fig. 1.**
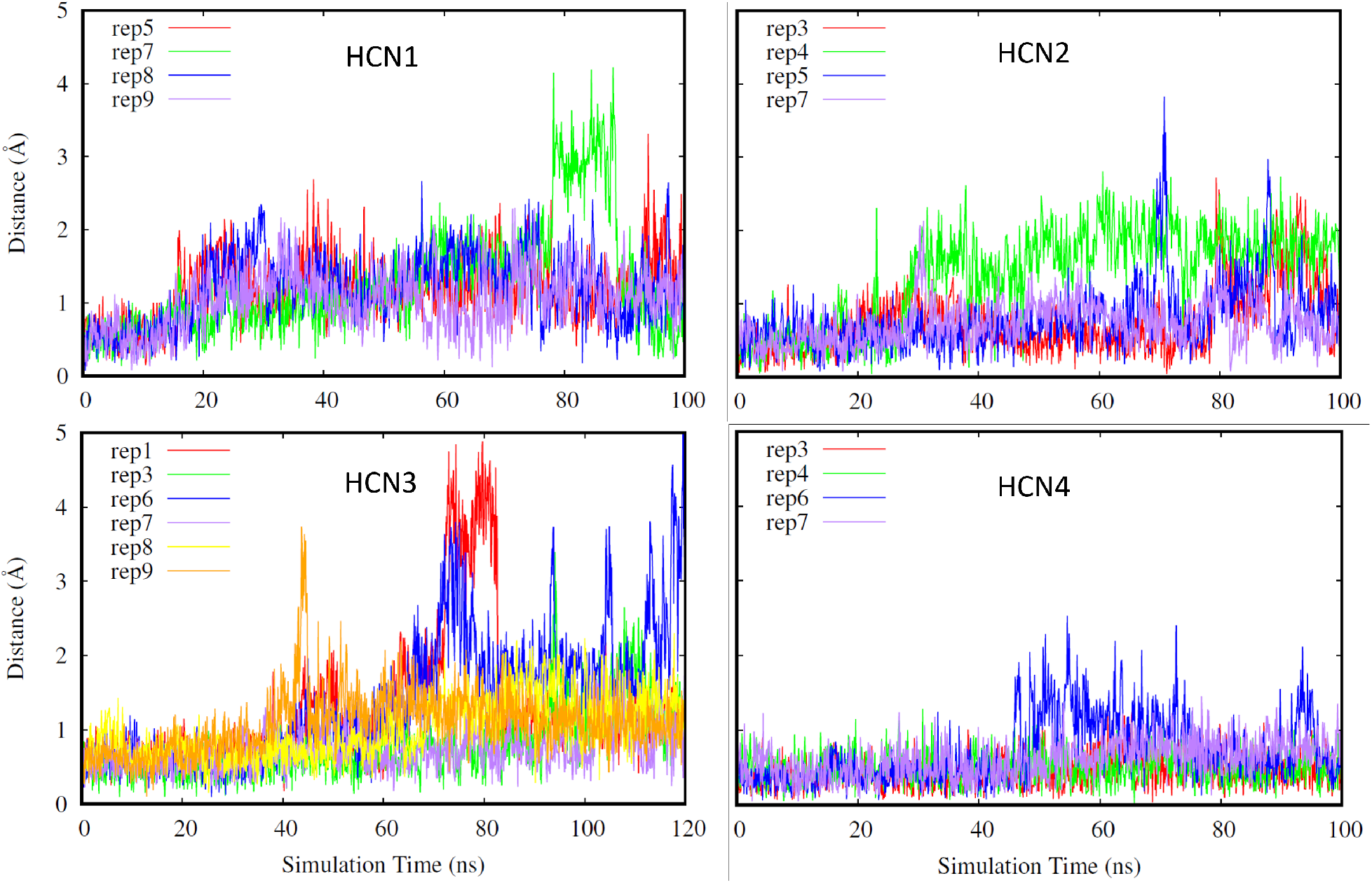
Ligand center of mass displacement for the replicates chosen for FEP for HCN isoforms (HCN1: top left, HCN2: top right. HCN3: bottom left, HCN4: bottom right). Different replicates for each isoform are shown in different colors.

Furthermore, for our second selection criteria, we looked into the RMSD of the ligand to further determine if a replicate is suitable for FEP. As shown in Figure 2, replicates were chosen for FEP if the RMSD of the ligand was low and stable, below 3 Å. Low RMSD values of the ligand indicate that it stable in the binding pocket, and maintains a conformation closely resembling that of the deposited structure. Although the ligand’s RMSD increases above the 3 Å threshold in some of the HCN3 replicates, for a majority of the presented systems the ligand RMSD remains at or below 3 Å. The highest ligand RMSD values are observed in HCN3 (Figure 2 bottom left) and further indicate that because the ligand was not in the original model and had to be placed using HCN1 as a reference, using more replicates is justified to ensure an accurate representation of this isoform’s behavior.

**Fig. 2.**
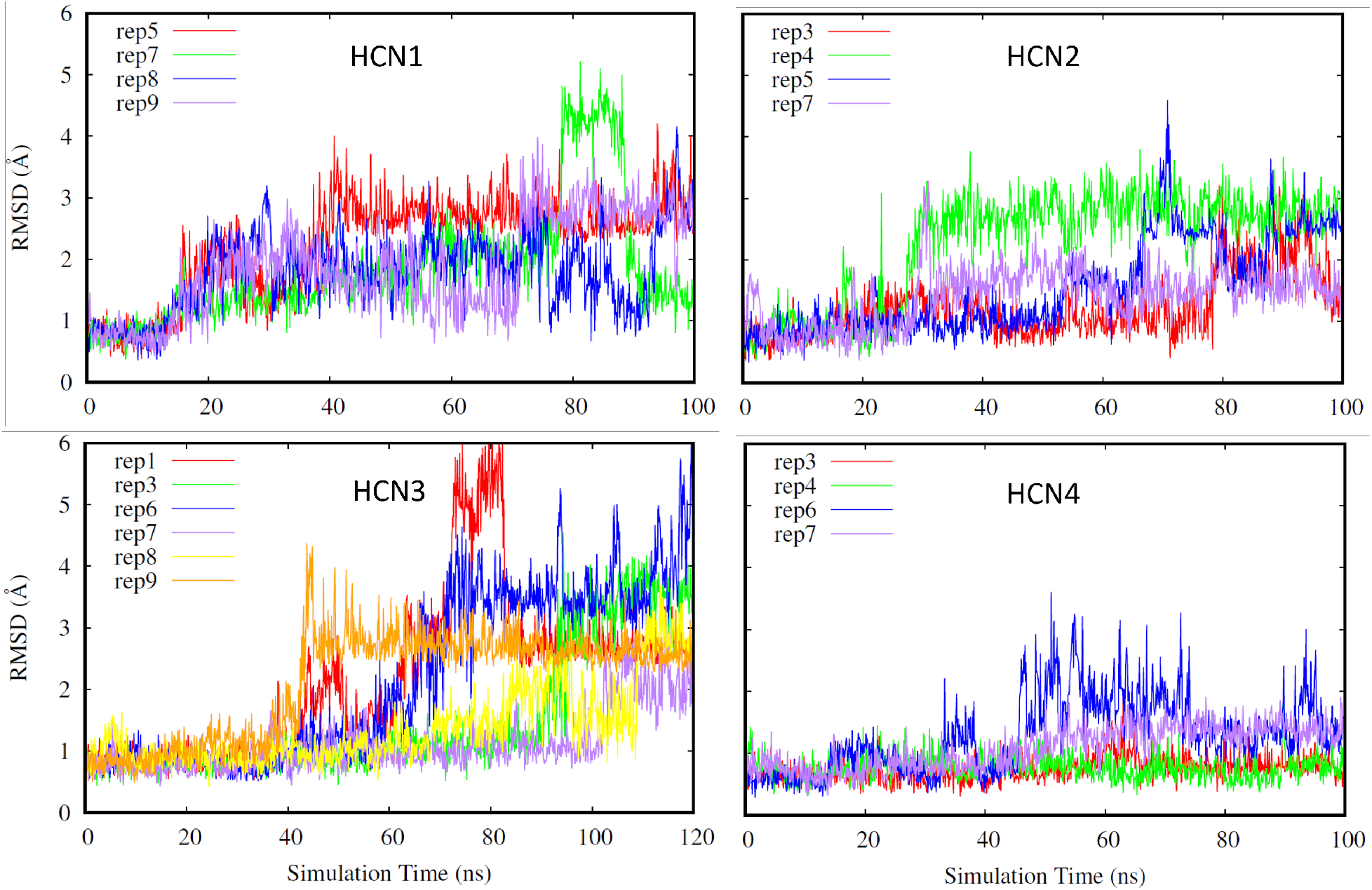
Ligand RMSD (HCN1: top left, HCN2: top right. HCN3: bottom left, HCN4: bottom right). Different replicates for each isoform are shown in different colors.

Additionally, we looked at the “lid distance” in Figure 3, defined as the distance between a particular residue in a given loop of the *β*-jelly roll and and a particular residue in the turn between helices C and D. A lower lid distance indicates that these two residues of the CNBD are in close proximity, meaning that the C helix is pulled in close to the CNBD and is barring the nucleotide inside the binding pocket much like a closed “lid”. While the ligand can continue to remain bound even if this “lid” should open, it can also be interpreted as another measure of domain stability and uniformity for it to remain closed. As seen in our previous analysis methods, the most variability is seen in HCN1 and HCN3, while the lid distance for HCN2 and HCN4 remains fairly stable throughout the simulations.

**Fig. 3.**
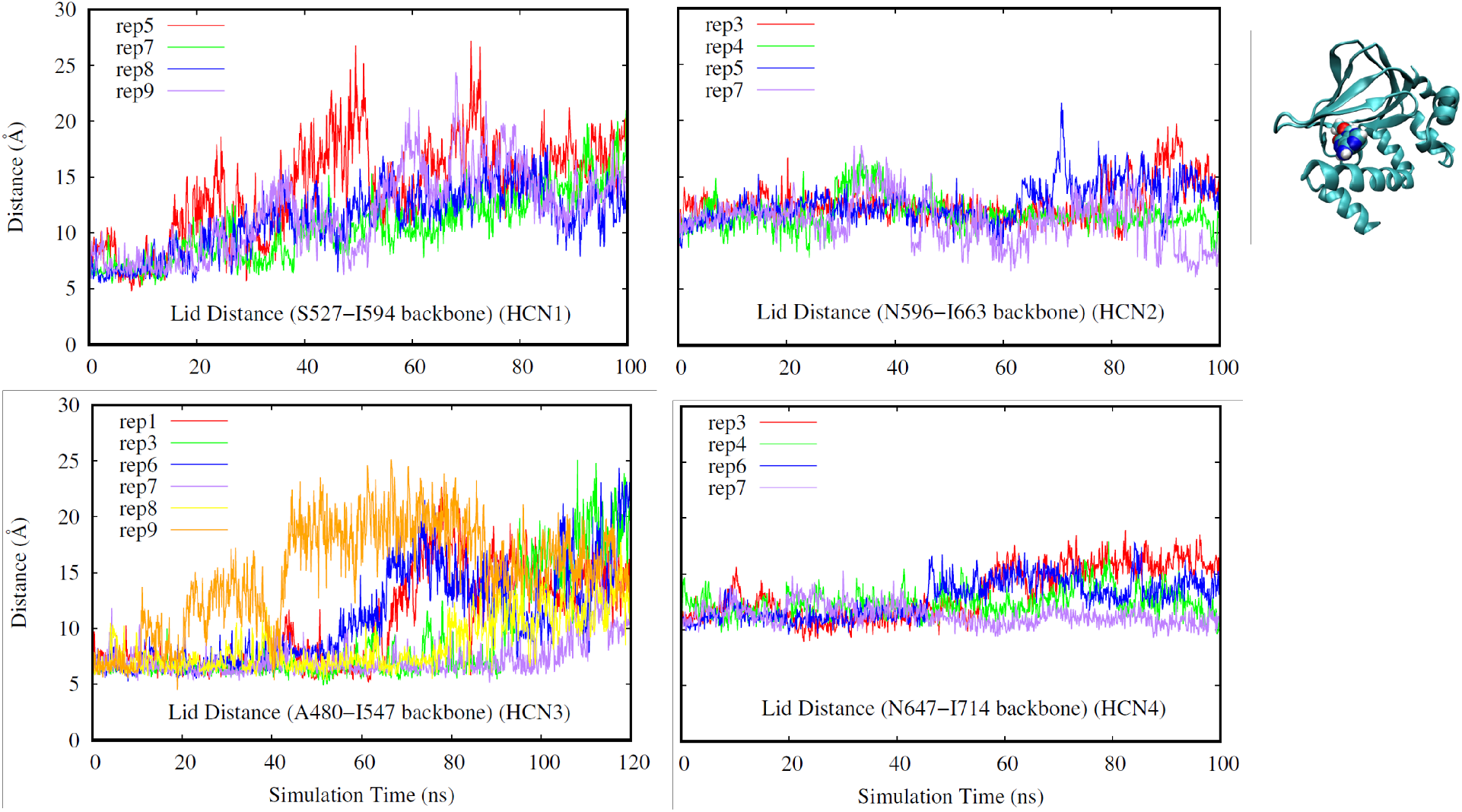
Lid Distance (HCN1: top left, HCN2: top right. HCN3: bottom left, HCN4: bottom right). Different replicates for each isoform are shown in different colors.

Additionally, we also considered the angle between helix B and helix C on the CNBD of HCN1-4 isoforms as shown in Figure 4. We chose to investigate this angle as another means to assess the behavior of the helix C “lid”, as well as overall domain stability. Although HCN2 and HCN4 show less fluctuation than HCN1 and HCN3 replicates, the angle between helices B and C remains fairly stable for all isoforms.

**Fig. 4.**
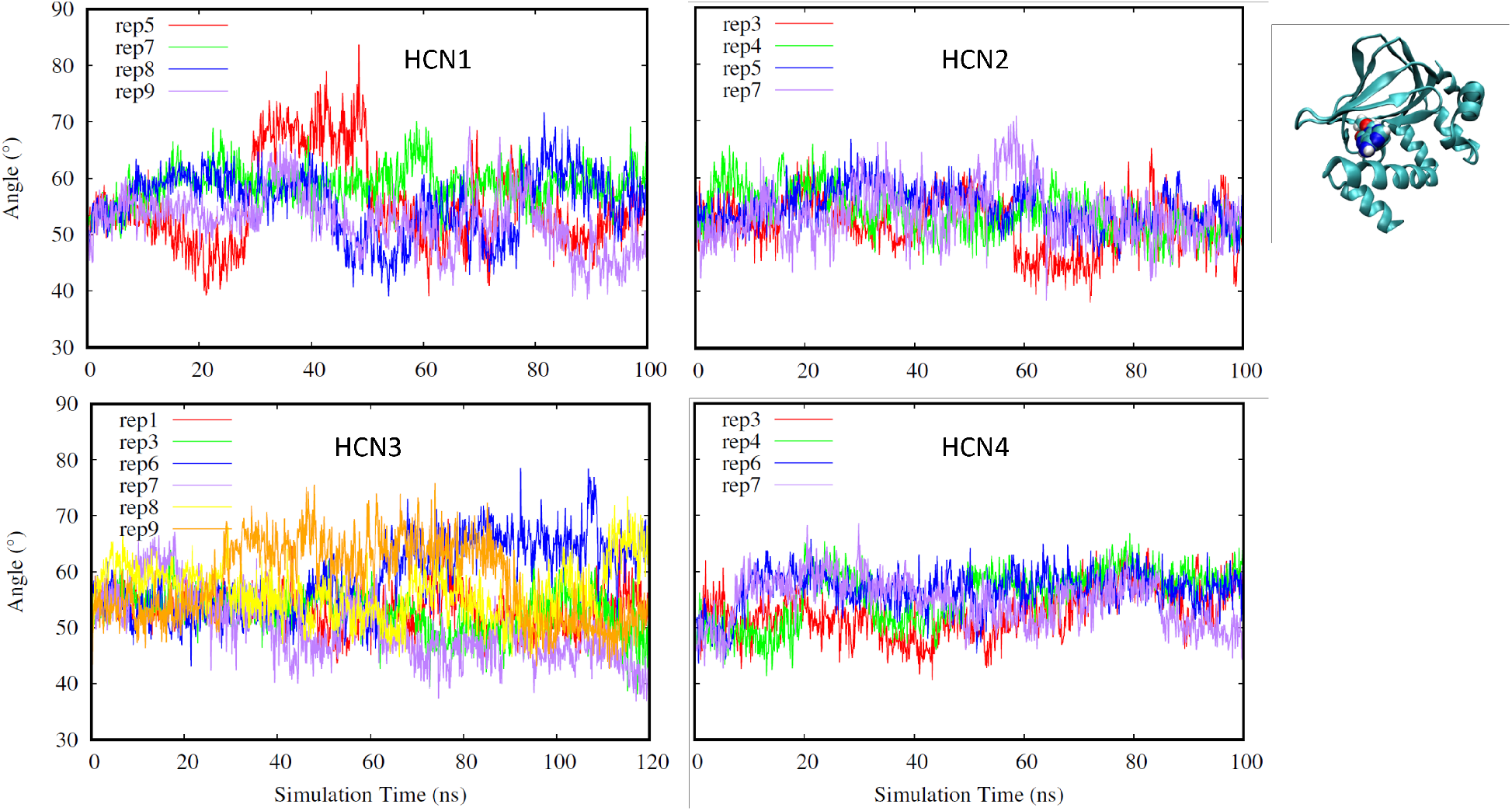
Angle between helices B and C (HCN1: top left, HCN2: top right. HCN3: bottom left, HCN4: bottom right). Different replicates for each isoform are shown in different colors.

Finally, for our last selection criteria we looked at hydrogen bonds that occur in the deepest part of the CNBD pocket. A replicate was chosen if the hydrogen bonds distances were low indicating stable bond formation as shown in Figures 5-7. Hydrogen bonds in the deepest part of the pocket indicate that the ligand is occupying the furthest point of the pocket away from the entry site, confirming that the ligand has firmly entered the binding region. Specifically, we have investigated a glycine, glutamate, and an arginine residue for all of the isoforms due to these residues being the deepest in the binding pocket.

**Fig. 5.**
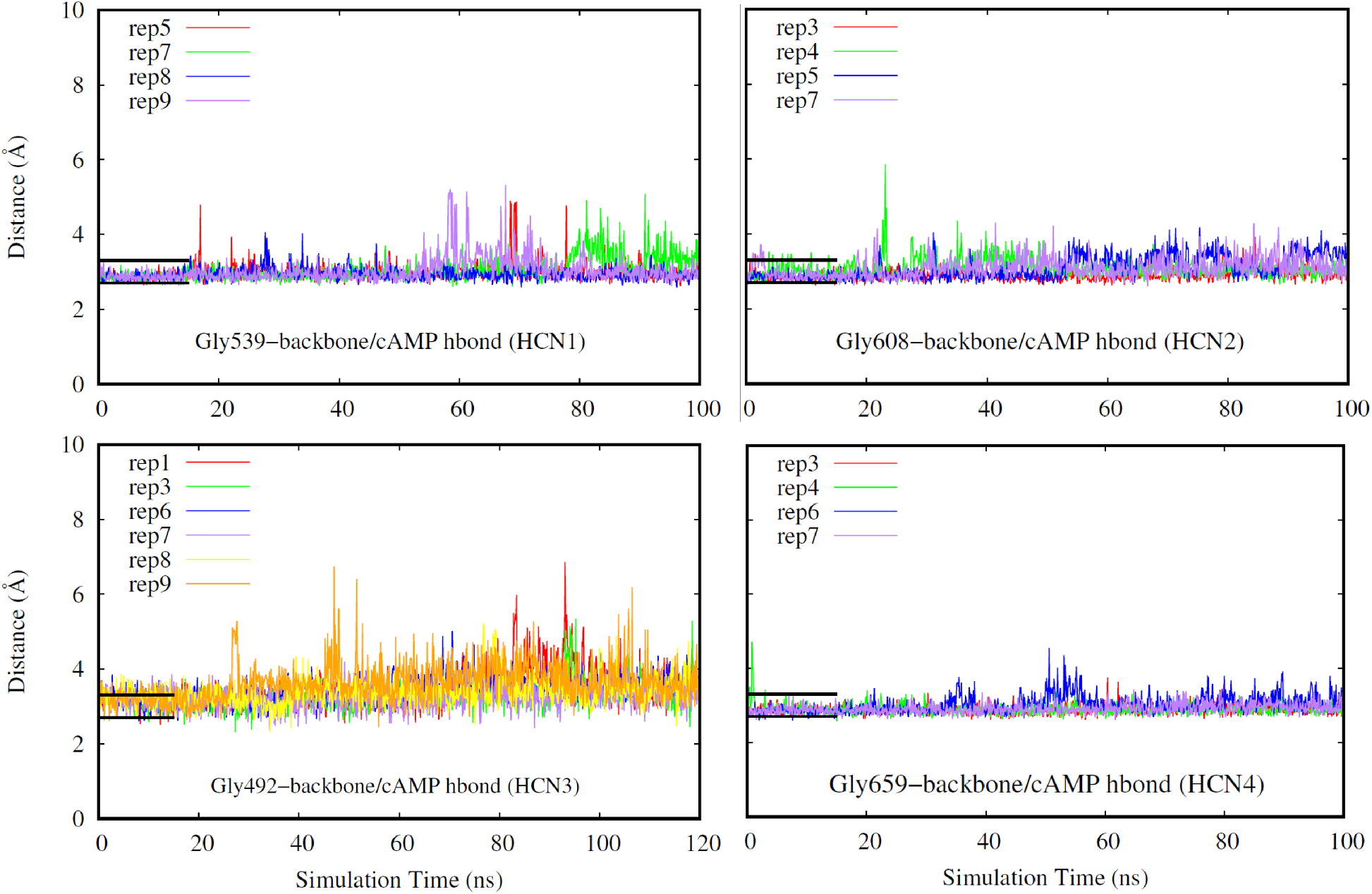
Hydrogen bond distance (HCN1: top left, HCN2: top right. HCN3: bottom left, HCN4: bottom right). Different replicates for each isoform are shown in different colors. Black bars indicate a range of distance values expected for typical hydrogen bonds, and their length corresponds to the duration the system was restrained.

**Fig. 6.**
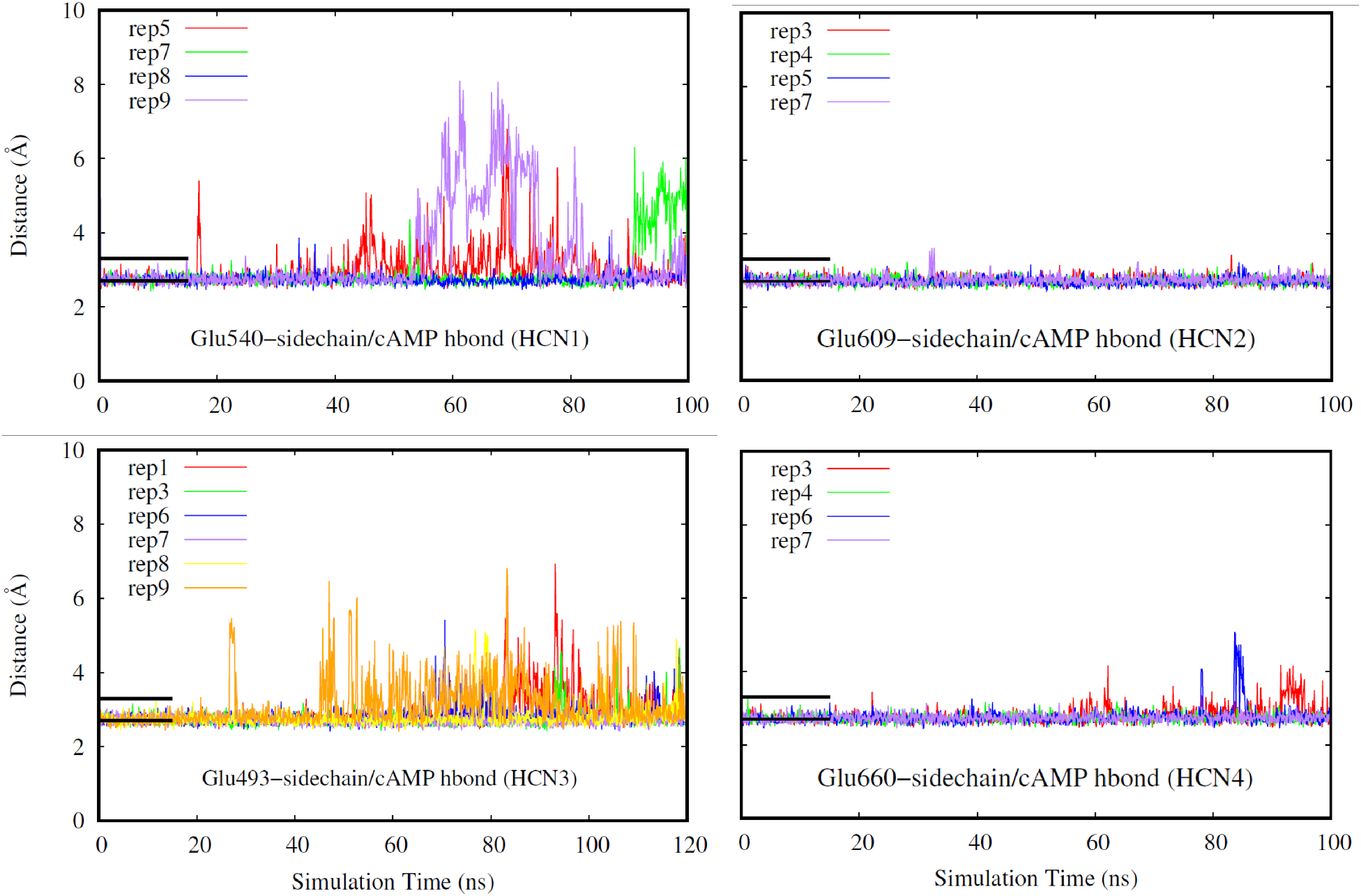
Hydrogen bond distance (HCN1: top left, HCN2: top right. HCN3: bottom left, HCN4: bottom right). Different replicates for each isoform are shown in different colors. Black bars indicate a range of distance values expected for typical hydrogen bonds, and their length corresponds to the duration the system was restrained.

**Fig. 7.**
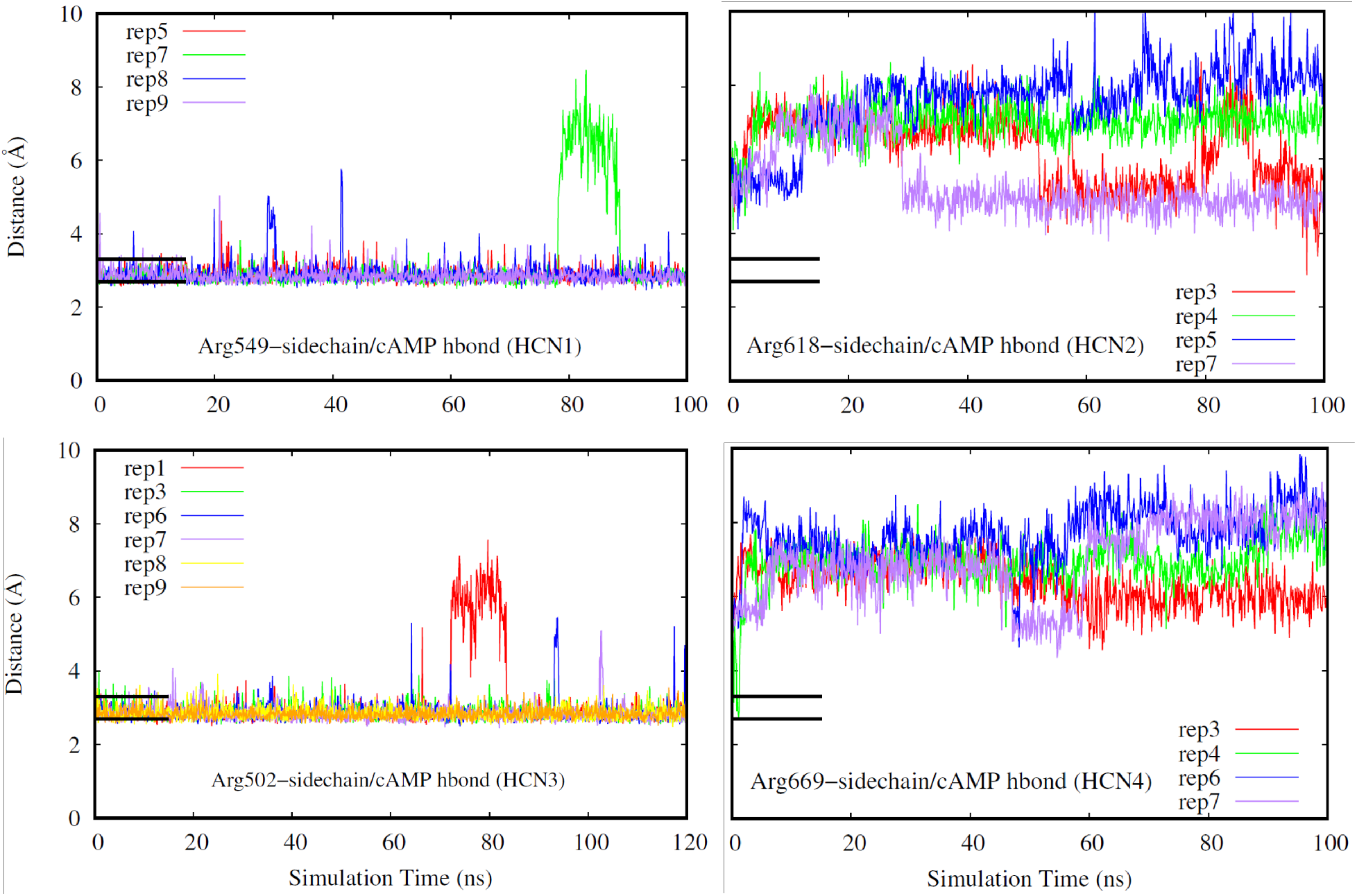
Hydrogen bond distance (HCN1: top left, HCN2: top right. HCN3: bottom left, HCN4: bottom right). Different replicates for each isoform are shown in different colors. Black bars indicate a range of distance values expected for typical hydrogen bonds, and their length corresponds to the duration the system was restrained.

Based on the replicates having low ligand center of mass displacement, low ligand RMSD, low lid distance values, a stable angle between helix B and helix C, and hydrogen bond formation in the deepest part of the CNBD pocket, we determined that replicates 5, 7, 8 and 9 should be used for FEP for HCN1. Similarly, for HCN2, we determined that replicates 3, 4, 5, and 7 will be used for FEP. For HCN3, we chose to use replicates 1, 3, 6, 7, 8, and 9 based on our aforementioned criteria. Additional replicates were used for HCN3 since our initial model was obtained from an AlphaFold model and the ligand was modeled using HCN1 as a reference. Finally, for HCN4, replicates 3, 4, 6, and 7 were chosen for FEP.

### FEP Reveals Differential Relative Binding Free-Energies for HCN Isoforms

It has been demonstrated previously that the HCN subfamily has different affinities for the binding of cyclic nucleotides, as well as differing sensitivities. ^10,12^ We used the FEP approach to estimate the binding free energy of cAMP bound to CNBDs of different HCN isoforms (see methods). The binding free energies were calculated using Δ*G*_−*solv*_(*cAMP*) = 118.63 ±0.39 kcal/mol (determined from a separate ligand-only simulation), and are shown in Table 1. Notably, these free energies possess a reasonable order of magnitude given the FEP method, and suggest that HCN1 and HCN3 have the highest comparable binding affinity, followed by HCN4, leaving HCN2 with the lowest binding affinity.

**Table 1:** Calculated FEP binding free energies of cAMP to Δ*G*_−*solv*_(*cAMP*) = 118.63 ±0.39 kcal/mol. System values reflect averages over 5 trajectories, with errors representing standard deviations.

| Isoform | Free Energy $\Delta G_{\text{protein}}$ (kcal/mol) | | | | | | $\Delta G_{\text{solv}} - \Delta G_{\text{protein}}$ (kcal/mol) |
| --- | --- | --- | --- | --- | --- | --- | --- |
|  | Pose 1 | Pose 2 | Pose 3 | Pose 4 | Pose 5 | Pose 6 |  |
| HCN1 | 129.415 ± 0.919 | 129.931 ± 1.485 | 128.421 ± 0.672 | 131.156 ± 0.660 | - | - | -11.097 ± 0.968 |
| HCN2 | 124.573 ± 1.826 | 123.230 ± 0.874 | 126.686 ± 1.170 | 127.466 ± 1.325 | - | - | -6.855 ± 1.115 |
| HCN3 | 129.415 ± 1.483 | 133.913 ± 2.762 | 130.151 ± 1.255 | 133.422 ± 1.087 | 127.208 ± 1.676 | 127.777 ± 0.654 | -11.68 ± 1.157 |
| HCN4 | 125.181 ± 1.642 | 127.407 ± 1.275 | 128.306 ± 2.181 | 129.107 ± 1.250 | - | - | -8.866 ± 1.209 |

### Per-Residue Interaction Free Energies Highlight the Importance of Conserved Arginine and Glutamate Residues

The FEP trajectories were used to determine the relative per-residue interaction free energies between the cyclic nucleotide and specific residues of interest using the equations described in the methods. The results of this analysis are shown in Table 2. All non-conserved residues with unobstructed access to the ligand, as well as any residues with demonstrable interactions such as hydrogen bonding, were considered for this investigation. Due to the high degree of sequence homology between the CNBDs, and to prevent confusion from comparing multiple chains with different sequence numbering, residues are referenced by their “residue offset”, defined as the sequence distance from the first considered residue. A full table listing the residue identities and offsets for each isoform is included at Table 3. The results show that HCN1 and HCN3 have the strongest interaction with the conserved arginine residue (offset +47). However, HCN2 and HCN4, which demonstrably lose this interaction in our simulations (Figure 7), seem to compensate by interacting more strongly with the conserved glutamate residue (offset +38). Our results align with the hydrogen bond occupancies and calculated binding free energies reported previously in this manuscript.

**Table 2:** Calculated per-residue interaction free energy contributions for CNBDs of HCN1-4.

| Reason for Consideration | Residue Offset | Relative Per-Residue Interaction Free Energy Contribution (kcal/mol) |  |  |  |
| --- | --- | --- | --- | --- | --- |
|  |  | HCN1 | HCN2 | HCN3 | HCN4 |
| non-conserved | +0 | -0.275 | -0.158 | -0.244 | -0.198 |
| non-conserved | +1 | 0.429 | 0.929 | 0.373 | 1.324 |
| non-conserved | +5 | 0.237 | 0.775 | 0.274 | 0.263 |
| non-conserved | +6 | 0.329 | -0.045 | 0.171 | -0.042 |
| non-conserved | +8 | -0.209 | -0.033 | -0.212 | 0.096 |
| non-conserved | +17 | 0.022 | 0.026 | 0.022 | 0.021 |
| non-conserved | +18 | -0.101 | -0.005 | 0.098 | 0.028 |
| non-conserved | +19 | -0.100 | -0.254 | -0.218 | -0.136 |
| non-conserved | +21 | -0.585 | -0.172 | -0.604 | -0.218 |
| non-conserved | +22 | -0.424 | -0.346 | 0.122 | -0.226 |
| non-conserved | +23 | 0.059 | -0.102 | 0.144 | -0.266 |
| non-conserved | +24 | -0.037 | 0.028 | 0.017 | -0.014 |
| non-conserved | +25 | -0.052 | -0.081 | 0.090 | 0.028 |
| non-conserved | +26 | 0.170 | 0.364 | 0.247 | -0.212 |
| non-conserved | +27 | -0.289 | -0.330 | -0.535 | -2.948 |
| non-conserved | +28 | 2.018 | 1.494 | 0.703 | 0.870 |
| non-conserved | +29 | 0.020 | 0.088 | -0.129 | -0.033 |
| non-conserved | +31 | 0.044 | 0.188 | 0.060 | 0.148 |
| h-bond | +37 | -2.360 | 0.214 | 1.904 | 1.450 |
| h-bond | +38 | -14.106 | -23.245 | -14.866 | -17.994 |
| h-bond | +40 | -1.362 | -0.294 | -1.836 | -0.036 |
| non-conserved | +44 | -0.384 | -0.126 | -0.442 | -0.140 |
| h-bond | +47 | -20.117 | -6.216 | -18.252 | -4.068 |
| h-bond | +48 | -1.774 | -3.108 | -2.462 | -2.831 |
| non-conserved | +86 | -0.131 | -0.218 | -0.203 | -0.162 |
| h-bond | +88 | 3.168 | 4.166 | 3.304 | 2.309 |
| non-conserved | +90 | -0.403 | -0.517 | -0.124 | -0.556 |

**Table 3:** Residue mapping across HCN1–4 isoforms based on residue offset.

| Offset | HCN1 | HCN2 | HCN3 | HCN4 |
| --- | --- | --- | --- | --- |
| +0 | I502 | I571 | V455 | I622 |
| +1 | I503 | I572 | V456 | I623 |
| +5 | A507 | T576 | S460 | T627 |
| +6 | V508 | I577 | V461 | I628 |
| +8 | K510 | K579 | R463 | K630 |
| +17 | V519 | V588 | L472 | V639 |
| +18 | A520 | V589 | L473 | V640 |
| +19 | G521 | S590 | S474 | S641 |
| +21 | I523 | L592 | L476 | L643 |
| +22 | T524 | T593 | A477 | T644 |
| +23 | K525 | K594 | R478 | K645 |
| +24 | S526 | G595 | G479 | G646 |
| +25 | S527 | N596 | A480 | N647 |
| +26 | K528 | K597 | R481 | K648 |
| +27 | E529 | E598 | D482 | E649 |
| +28 | M530 | M599 | T483 | T650 |
| +29 | K531 | K600 | R484 | K651 |
| +31 | T533 | S602 | T486 | A653 |
| +37 | G539 | G608 | G492 | G659 |
| +38 | E540 | E609 | E493 | E660 |
| +40 | C542 | C611 | C495 | C662 |
| +44 | K546 | R615 | R499 | R666 |
| +47 | R549 | R618 | R502 | R669 |
| +48 | T550 | T619 | T503 | T670 |
| +86 | I588 | I657 | M541 | L708 |
| +88 | R590 | R659 | R543 | R710 |
| +90 | D592 | D661 | L545 | D712 |

## Conclusion

In conclusion, our comparative MD simulations offer insights into the binding of cAMP in HCN channels, and attempts to establish a molecular basis for the difference in sensitivities between isoforms. Our FEP results suggest that HCN1 and HCN3 have the highest comparable binding affinity, followed by HCN4, leaving HCN2 with the lowest relative binding affinity compared to the other isoforms. Additionally, the importance of the conserved arginine residue (offset +47) in HCN1 and HCN3 as well as the conserved glutamate residue (offset +39) in HCN2 and HCN4 within the CNBD were highlighted by showing that these residues have the highest interaction free-energy with the cAMP ligand. Our results from the per-residue interaction free-energy calculations are also consistent with our observations regarding the hydrogen bonds that form in the deepest parts of the CNBD pocket.

Collectively, our findings shed light on the specific binding interactions that drive channel modulation, as well as the factors that contribute to differences in channel sensitivity to cAMP across HCN isoforms. These insights offer valuable context for progressing the understanding of HCN channel modulation by cAMP, which could be used to develop selective drugs targeting isoforms of the channel.

## Supplemental Information Available

Figures and information in the Supplemental Information provide additional analysis for the replicates of the HCN isoforms that were not chosen for the FEP approach based on our MD simulations as discussed in the manuscript.

## Acknowledgement

This research was supported by the National Institute of General Medical Sciences (NIH grant R35GM147423 awarded to M.M.), the National Science Foundation (NSF grant CHE 1945465 awarded to M.M.), and the Arkansas Biosciences Institute. Computational resources were provided by the Texas Advanced Computing Center (TACC) at the University of Texas at Austin (Frontera) through LRAC allocation CHE21003 to M.M. The work also used Stampede at TACC and Bridges-2 at the Pittsburgh Supercomputing Center through allocation MCB150129 from the Advanced Cyberinfrastructure Coordination Ecosystem: Services & Support (ACCESS) program. Additional computational support came from the Arkansas High-Performance Computing Center, funded by multiple NSF grants and the Arkansas Economic Development Commission. We also thank Dr. Adithya Polasa for helpful discussions and continuous support.

