## Supporting Information for "Predicting Binding Affinities for the Binding Domain of Hyperpolarization-Activated Cyclic Nucleotide-Gated Channel Isoforms Using Free-Energy Perturbation"

Before conducting FEP to obtain absolute binding free-energies for cAMP binding to the CNBD of isoforms of HCN, we first identified suitable replicates from our equilibrium simulations to conduct FEP on for each isoform. Specific replicates were chosen to ensure that the ligand remained suitably bound once the restraints were lifted in preparation for the FEP step. In the following section, we discuss our selection criteria for rejecting certain replicates to conduct FEP on for each isoform, and present data for these rejected trajectories. In general, replicates were chosen if they had low ligand center of mass displacement, low ligand RMSD, and stable hydrogen bonds with key residues of interest. Replicates that did not meet these criteria were deemed to be unsuitable for providing accurate binding results for FEP.

For our first selection criteria, we targeted replicants whose ligand center of mass displacements remained stable and did not exceed 2 Å. A small ligand center of mass displacement indicates that the ligand was not changing its overall position relative to the binding pocket, ensuring that the ligand was not in the process of leaving the pocket. With this in mind, replicates were not chosen to advance to the FEP stage if the ligand center of mass displacement was above 2 Å, as this indicates that the ligand was changing its overall position in the binding pocket and was not stably bound (Fig. 8). If we were to run FEP on the ligand when it is not stably bound in the pocket, our free-energy calculations would be skewed. Therefore, to avoid the consideration of large, irrelevant fluctuations, we have excluded certain replicates of the isoforms to conduct the FEP approach on.

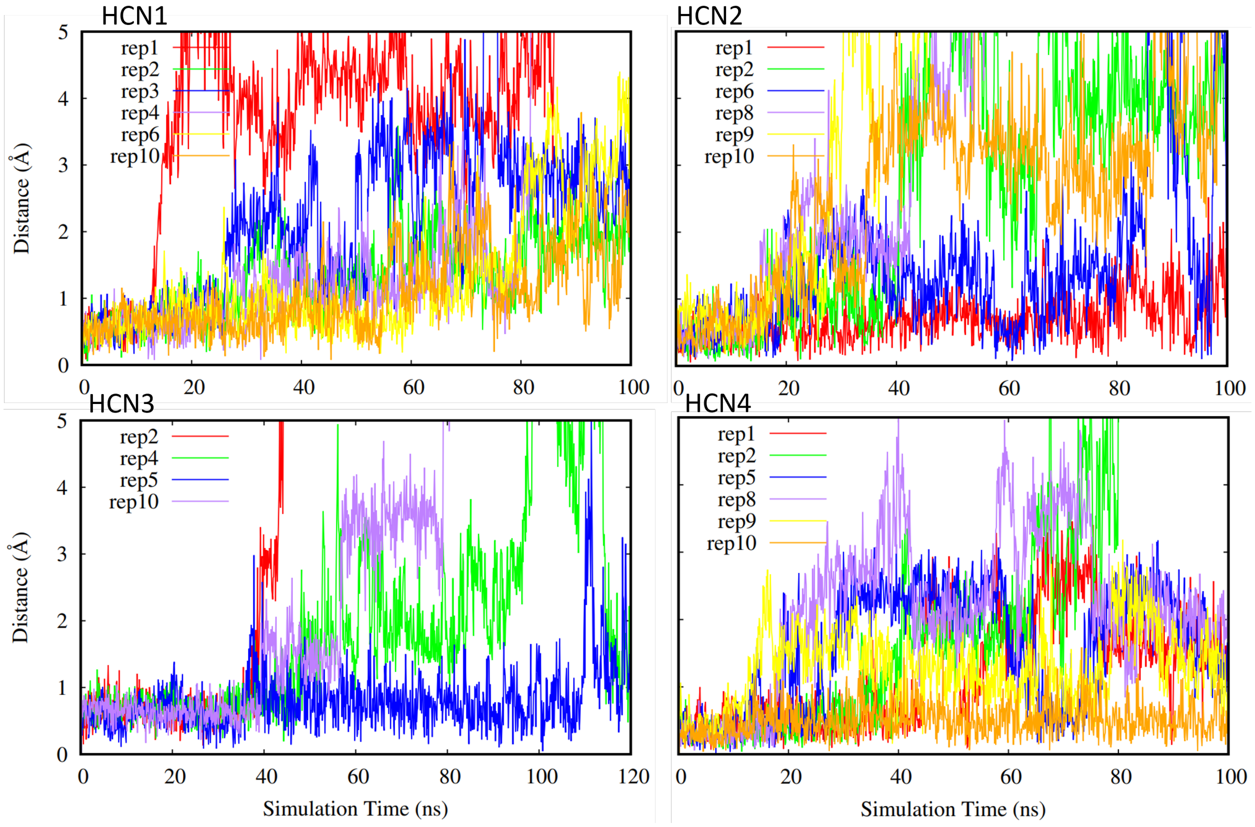


**Fig. S1.** Ligand center of mass displacement for the replicates not chosen for FEP for HCN isoforms (HCN1: top left, HCN2: top right. HCN3: bottom left, HCN4: bottom right). Different replicates for each isoform are shown in different colors.

For our second selection criteria, we used the RMSD of the ligand in much the same way as the ligand center of mass. Replicates were chosen for FEP if the RMSD of the ligand remained stable and below 3 Å. Low RMSD values of the ligand indicate that it is stable in the binding pocket, and maintains a conformation closely resembling that of the deposited structure. For a similar reason as the ligand center of mass displacement criteria, replicates were not chosen for the FEP stage if they displayed ligand RMSD values higher than 3 Å, as this indicates potential positional and conformation instability (Fig. 9).

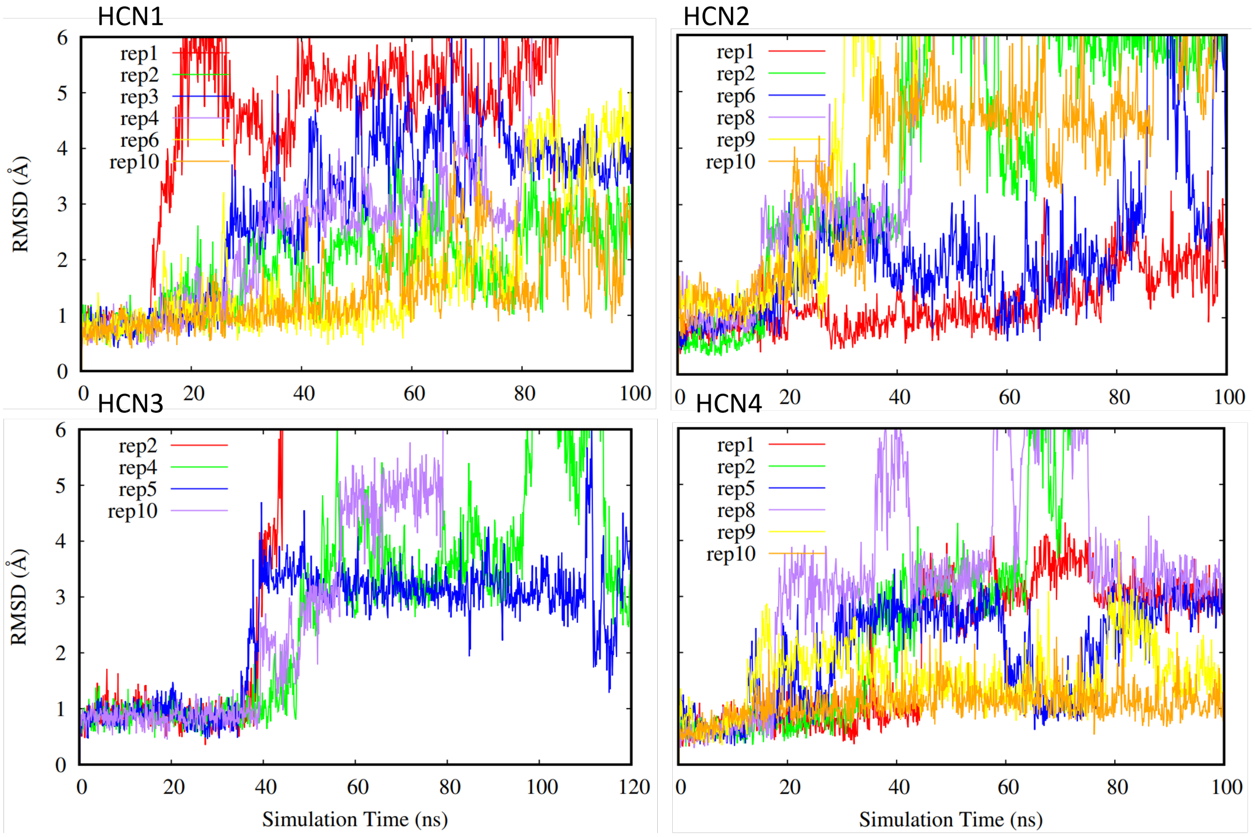


**Fig. S2.** Ligand RMSD (HCN1: top left, HCN2: top right. HCN3: bottom left, HCN4: bottom right). Different replicates for each isoform are shown in different colors.

Additionally, we looked at the "lid distance" to determine if a replicate was suitable for FEP. A high lid distance indicates that residues of the CNBD are not in close proximity, meaning that the C helix is pulled away from the $\beta$-jelly roll, thus allowing the nucleotide to exit the binding pocket much like a open "lid". While the ligand could continue to remain bound even if this "lid" should open, it can also be interpreted as another measure of domain stability and uniformity for it to remain closed. Therefore, replicates were not chosen if the C-helix was not close in proximity to the $\beta$-jelly roll, as this would imply that the CNBD was open (Fig. 10).

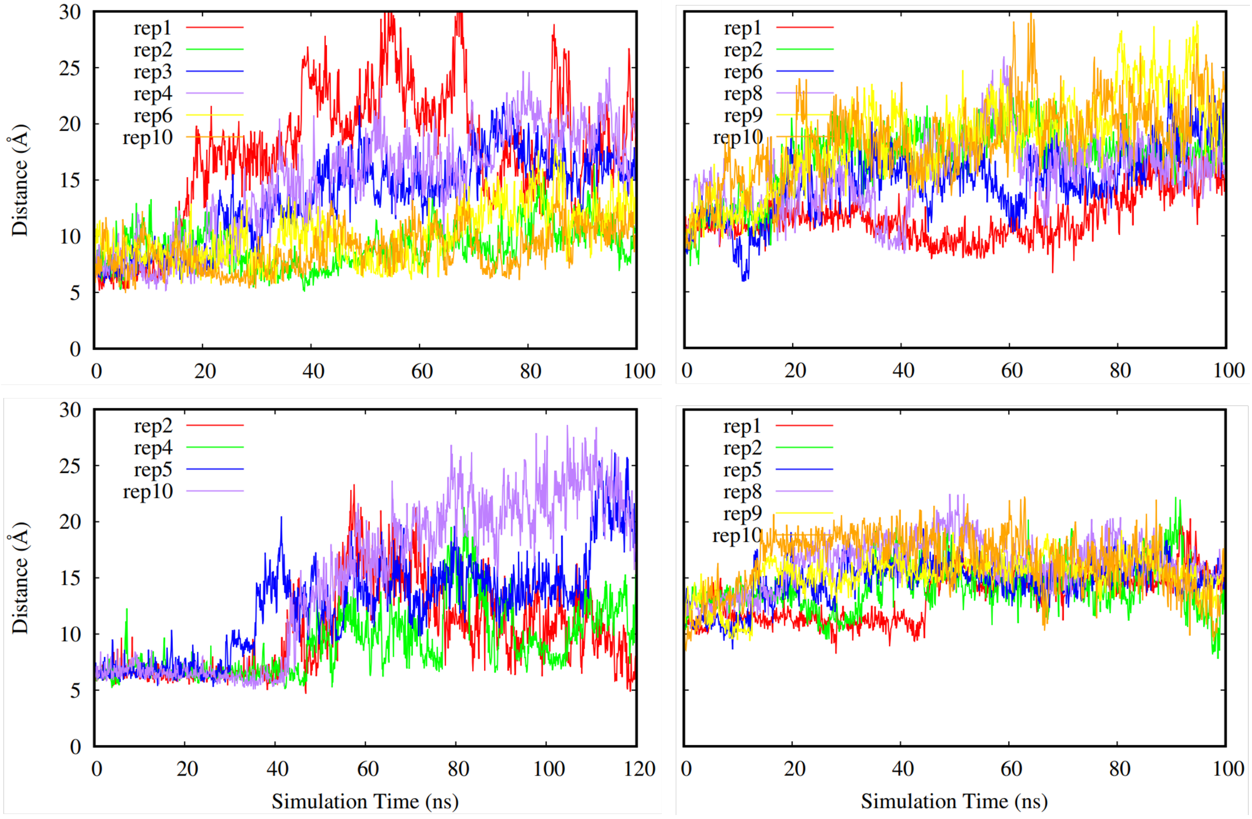


**Fig. S3.** Lid Distance (HCN1: top left, HCN2: top right. HCN3: bottom left, HCN4: bottom right). Different replicates for each isoform are shown in different colors.

Additionally, we also considered the angle between helix B and helix C on the CNBD of HCN1-4 isoforms. We chose to investigate this angle as another means to assess the behavior of the helix C ”lid”, as well as overall domain stability. For a majority of the replicates, this angle remains relatively stable. However, there are more fluctuations in the angles for the replicates that were not chosen for FEP than the ones that were chosen (Fig. 11).

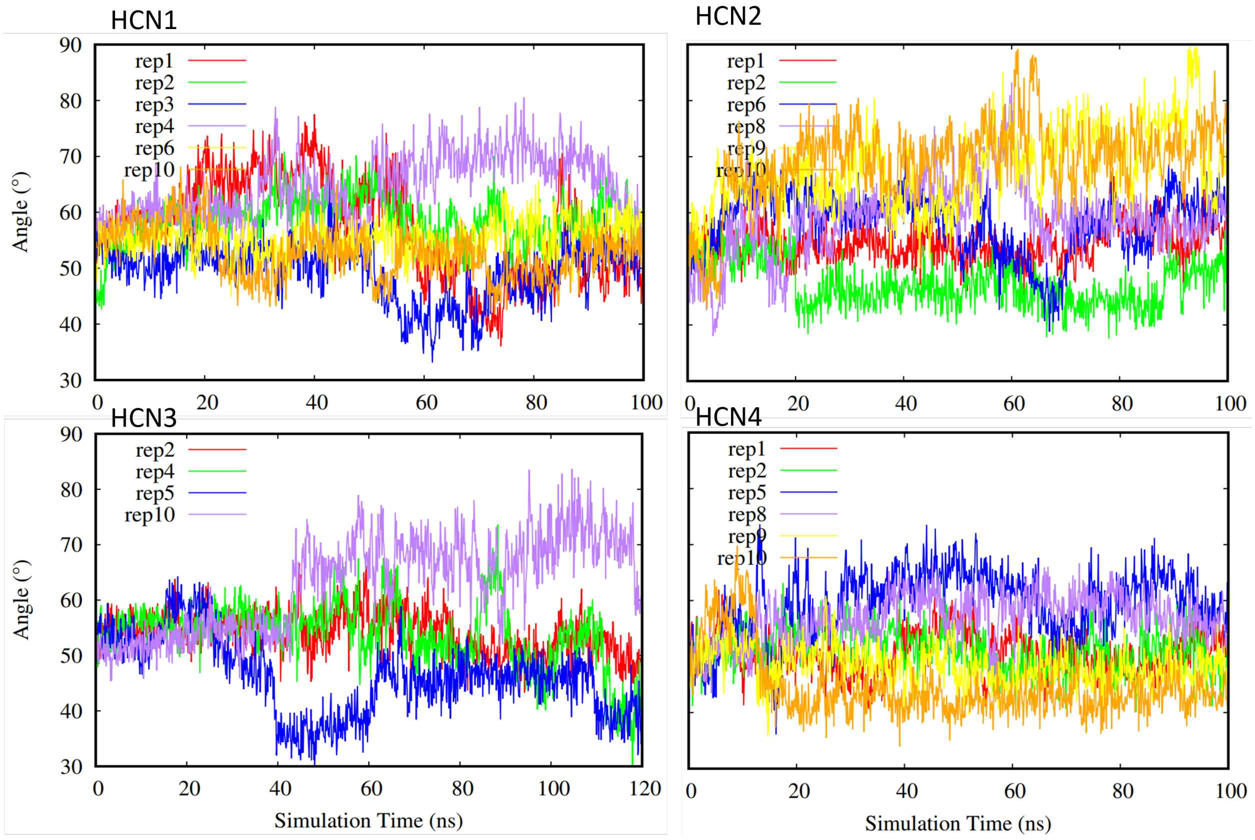


**Fig. S4.** Angle between helices B and C (HCN1: top left, HCN2: top right. HCN3: bottom left, HCN4: bottom right). Different replicates for each isoform are shown in different colors.

Finally, for our last selection criteria we looked at hydrogen bonds that occur in the deepest part of the CNBD pocket. A replicate was not chosen if the hydrogen bonds distances were high indicating stable bond formation did not occur as shown in Figures 12-14. Hydrogen bonds in the deepest part of the pocket indicate that the ligand is occupying the furthest point of the pocket away from the entry site, confirming that the ligand has firmly entered the binding region. Specifically, we have investigated a glycine, glutamate, and an arginine residue for all of the isoforms due to these residues being the deepest in the binding pocket. Without these bonds present, the ligand cannot occupy the full binding pocket.

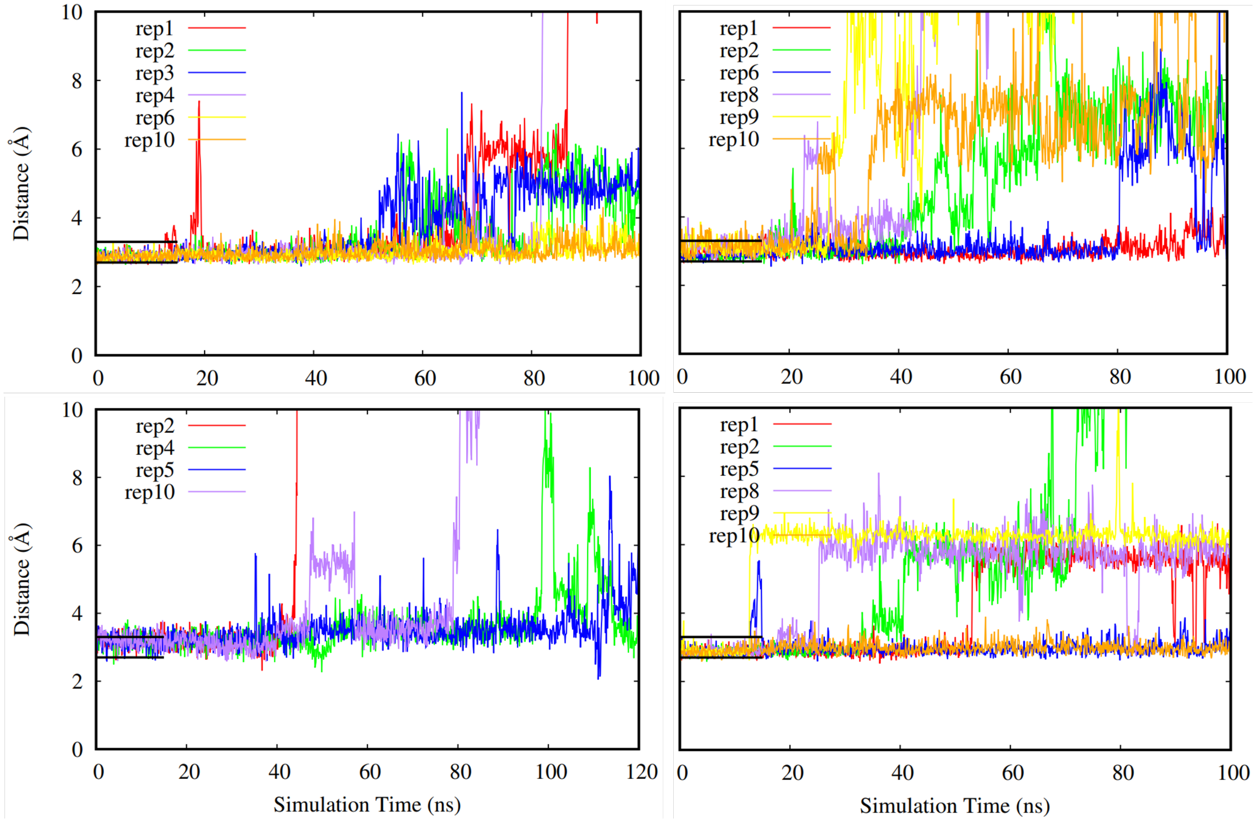


**Fig. S5.** Hydrogen bond distance (HCN1: top left, HCN2: top right. HCN3: bottom left, HCN4: bottom right). Different replicates for each isoform are shown in different colors. Black lines on the plot indicate the distance values indicative of a hydrogen bond. The lengths of the black lines are the length of time that the restraints were in place.

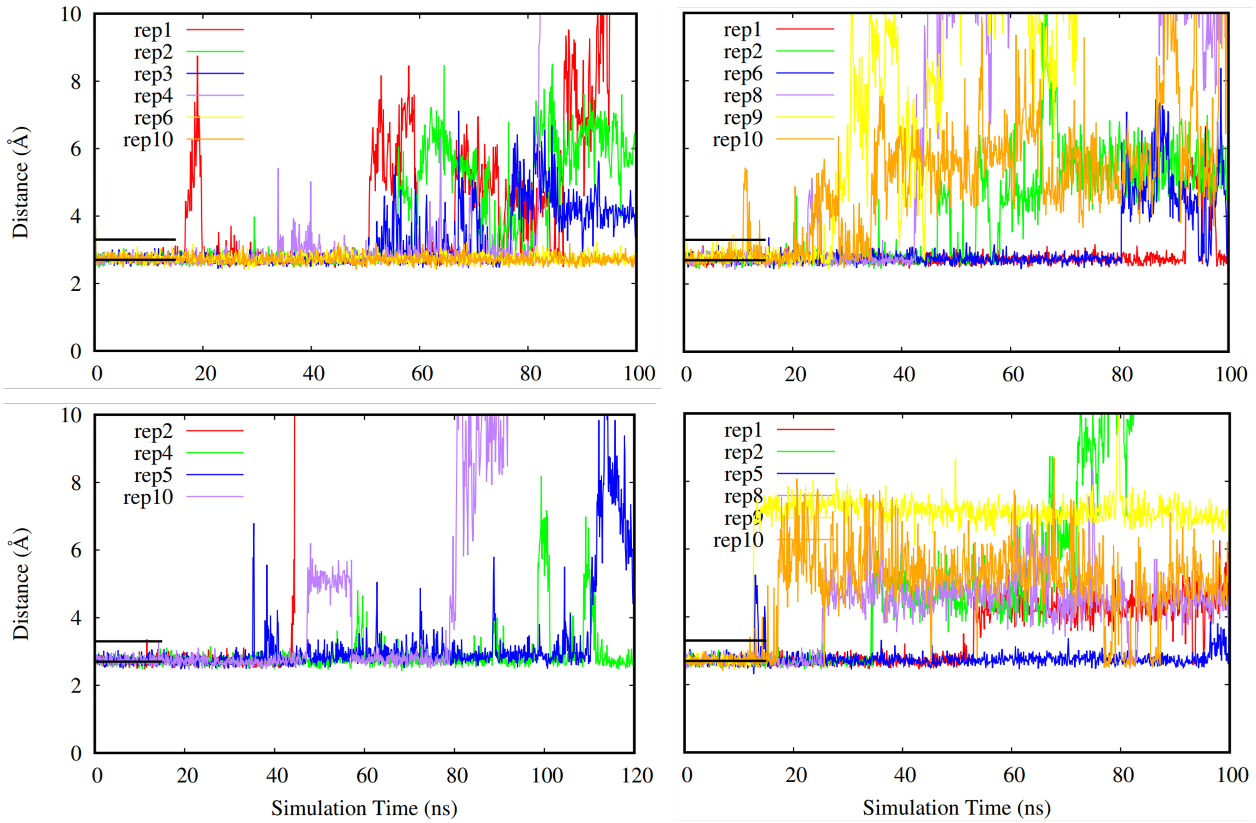


**Fig. S6.** Hydrogen bond distance (HCN1: top left, HCN2: top right. HCN3: bottom left, HCN4: bottom right). Different replicates for each isoform are shown in different colors. Black lines on the plot indicate the distance values indicative of a hydrogen bond. The lengths of the black lines are the length of time that the restraints were in place.

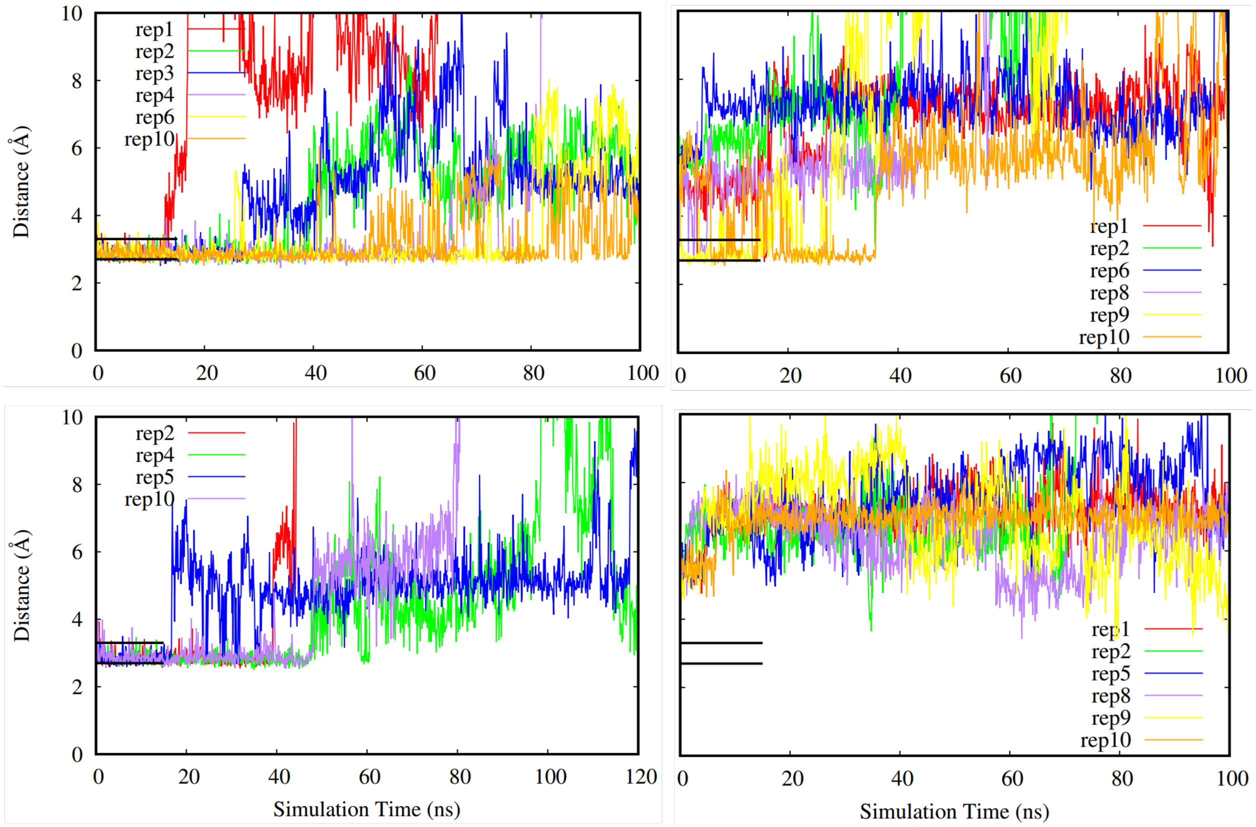


**Fig. S7.** Hydrogen bond distance (HCN1: top left, HCN2: top right. HCN3: bottom left, HCN4: bottom right). Different replicates for each isoform are shown in different colors. Black lines on the plot indicate the distance values indicative of a hydrogen bond. The lengths of the black lines are the length of time that the restraints were in place.

Based on the replicates having high ligand center of mass displacement, high ligand RMSD, high lid distance values, an unstable angle between helix B and helix C, and unstable hydrogen bond formation in the deepest part of the CNBD pocket, we determined that replicates 1, 2, 3, 4 and 10 should not be used for FEP for HCN1. Similarly, for HCN2, we determined that replicates 1, 2, 6, 8, 9, and 10 should not be used for FEP. For HCN3, we chose to not use replicates 2, 4, 5, and 10 based on our aforementioned criteria. Finally, for HCN4, replicates 1, 2, 5, 8, 9, and 10 were not chosen for FEP.
